# Harnessing A Unified Multi-modal Sequence Modeling to unveil Protein-DNA Interdependency

**DOI:** 10.1101/2025.02.26.640480

**Authors:** Mingchen Li, Yuchen Ren, Peng Ye, Jiabei Cheng, Xinzhu Ma, Yuchen Cai, Wanli Ouyang, Bozitao Zhong, Banghao Wu, Nanqing Dong, Liang Hong, Pan Tan

## Abstract

The interdependent orchestration of proteins, coding DNA sequences (CDS), and adjacent regulatory DNA elements underpins biological functionality, where evolutionary constraints shape codon usage, transcriptional regulation, and structural stability. We present ProDMM, a multimodal framework that deciphers these systemic relationships through unified sequence modeling. Integrating a context-aware encoder with conditional generation, our architecture captures codependent patterns between protein motifs and cis-regulatory syntax. ProDMM’s encoder demonstrates superior predictive capabilities in forecasting pathway-level phenotypes and protein expression profiles, outperforming conventional single-modality approaches. Its decoder generates synthetically optimized regulatory elements by resolving spatial dependencies between coding sequences and adjacent non-coding regions, enabling transcriptional fine-tuning unavailable to sequence-agnostic methods. By resolving how localized CDS-NCDS-protein interdependencies propagate to system-level phenotypes, ProDMM establishes a paradigm for holistic biological sequence engineering. This approach enables coordinated optimization of translation efficiency, enzymatic activity, and pathway flux, offering transformative potential for designing tailored biosynthesis systems and advancing sustainable bioproduction strategies.

## 1 Introduction

Synthetic biology is at the forefront of biological engineering, enabling the systematic redesign of living systems through molecular component assembly. This discipline merges the precision of protein engineering in the creation of functional biomolecules with the capacity of metabolic engineering for the orchestration of pathways, enabling groundbreaking applications from precision therapeutics to renewable biomanufacturing [1]. However, a fundamental challenge persists: the interdependent treatment of protein coding sequences (CDS), their translated products, and adjacent noncoding regulatory elements (NCDS) that collectively govern biological outcomes. While codon selection influences translational efficiency, promoter architectures dictate transcriptional initiation, and protein folding dictates functional integrity, conventional approaches addressing these elements in isolation systematically fail to capture their evolutionary synchrony—a limitation starkly evident in nature’s own designs. The limitations of conventional methodologies become starkly apparent when examining nature’s own engineering solutions. In extremophile organisms, thermal adaptation manifests itself through coordinated optimization of protein thermostability and promoter region GC content, a dual strategy in which heat-resistant enzymes pair with thermally stable DNA structures [2]. Similarly, high-expression bacterial operons exhibit precise alignment between upstream regulatory elements and codon patterns [3]. These principles of biological design reveal a fundamental truth: optimal functionality emerges not from isolated sequence optimization but from the harmonious integration of coding and regulatory elements. Traditional approaches that separately engineer promoters, codon usage, and protein sequences inevitably disrupt these evolved relationships, leading to suboptimal expression and metabolic imbalances [4].

Recent breakthroughs in deep learning have revolutionized single-modality sequence analysis. Protein language models (e.g., ESM-2 [5], ProtTrans [6]) achieve atomic-level structure prediction through unsupervised learning on millions of sequences, while DNA foundation models (e.g., Evo [7], DNABERT-2 [8]) decode regulatory syntax across genomes. However, these single-modal approaches inherit the core weakness of the reductionist paradigm—they process proteins and DNA as separate linguistic domains, blind to the evolutionary constraints that bind them. A protein language model might perfectly predict enzyme thermostability, yet remains oblivious to the promoter adaptations that ensure its proper expression under high temperatures. Conversely, DNA models could predict promoter strength while might inadvertently selecting codon combinations that impede translation without the condition of protein information. This modality isolation creates artificial divides where nature maintains continuous dialogues, fundamentally constraining their utility in synthetic biology applications.

To bridge this divide, we introduce ProDMM (Protein-DNA Multimodal Model), a framework that redefines biological sequence analysis through unified representation of proteins, CDS, and adjacent regulatory contexts. ProDMM has been pre-trained in a self-supervised manner on over 129 million paired sequences, allowing it to integrate the semantic features of protein sequences, coding DNA sequences (CDS), and their adjacent non-coding DNA sequences (NCDS), while capturing their inter-dependencies. ProDMM’s architecture embodies a unified approach to biological sequence modeling through synergistic integration of a bidirectional encoder and conditional decoder. The encoder employs masked language modeling (MLM) to reconstruct evolutionary narratives from fragmented sequence data, establishing contextual relationships between protein structural motifs, their coding DNA sequences (CDS), and adjacent non-coding regulatory elements (NCDS). Complementing this, the decoder extends this biological intelligence through two key conditional generation tasks: protein-to-CDS reverse translation and CDS-to-NCDS synthesis preserving spatial constraints of native operons. Through autoregressive generation guided by evolutionary alignment, the decoder produces functional sequences where synthetic promoters inherently complement their downstream coding regions.

ProDMM’s capabilities are validated through comprehensive benchmarking against leading protein- and DNA-specific models. The framework demonstrates superior performance in zero-shot prediction of mutation fitness landscapes, transcriptional compatibility assessment, and metabolic pathway yield forecasting. Its conditional generation module produces synthetic promoters with enhanced functional synergy to target coding regions compared to sequence-agnostic approaches, while cross-modal representation learning enables accurate prediction of pathway-level behaviors from unlabeled sequence data. These advances establish a new paradigm for biological engineering—where machine learning transcends predictive modeling to become a discovery engine for evolutionary design principles. By preserving natural optimization strategies while enabling novel combinatorial exploration, ProDMM bridges the gap between nature’s wisdom and engineering ambition, offering transformative potential for sustainable bioproduction and precision bioengineering.

## 2 Results

### 2.1 Architecture of ProDMM

ProDMM employs a dual-component architecture built upon the Transformer framework [9], comprising two specialized sub-models: a multimodal encoder (ProDMM Encoder) and a conditional sequence generator (ProDMM Seq2Seq). As illustrated in Figure 1A, the encoder adopts a BERT-style [10] pretraining paradigm using MLM to learn contextual relationships across protein, coding DNA (CDS), and non-coding regulatory sequences. Additionally, the ProDMM Seq2Seq, shown in Figure 1B, extends this foundation by integrating the encoder with a task-specific decoder through cross-attention mechanisms, enabling autoregressive generation of biologically constrained sequences. Both components undergo hierarchical pretraining—first through self-supervised MLM on 129 million curated CDS-protein-regulatory element triplets from the Genome Taxonomy Database (GTDB), followed by conditional generation tasks that enforce evolutionary consistency between synthesized elements and their biological contexts. Once the pre-training phase is completed, ProDMM can be utilized in various scenarios and tasks, as demonstrated in 1C. The pretraining corpus, rigorously processed as detailed in Methods, captures comprehensive prokaryotic sequence diversity, with CDS lengths predominantly distributed between 200-400 nucleotides (Fig. S1A) and taxonomical representation spanning 23 bacterial phyla (Fig. S1B). This multimodal training strategy enables the model to establish three critical biological associations: (1) codon-to-amino acid mapping constraints, (2) promoter-CDS spatial relationships, and (3) protein-stability-guided regulatory element evolution. The architecture with joint representation learning enhances prediction accuracy for both molecular properties and system-level phenotypes.

**Figure 1:**
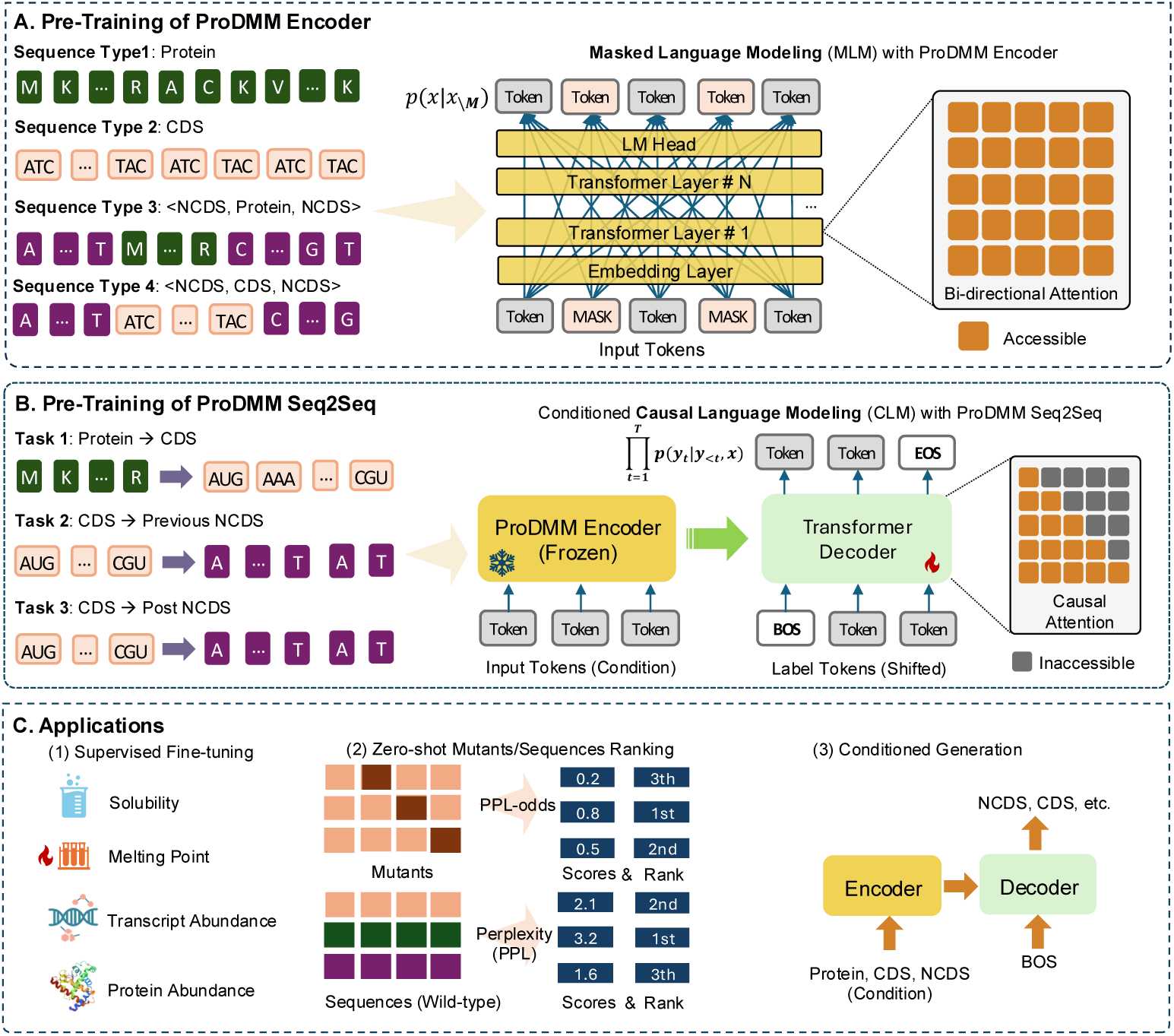
The architecture of ProDMM: (A) The ProDMM Encoder, a Transformer encoder, was pre-trained using masked language modeling on four types of sequences. (B) The ProDMM Seq2Seq equips the ProDMM Encoder with an additional decoder and was pre-trained with conditioned causal language modeling across three generation tasks. (C) Applications of ProDMM. It can be used as a pre-trained model for supervised fine-tuning across various tasks. It also functions as a model for scoring and ranking sequences or mutants (ProDMM Encoder) and for conditioned sequence generation (ProDMM Seq2Seq).

#### 2.1.1 ProDMM Encoder

ProDMM’s encoder architecture leverages a Transformer-based bidirectional attention mechanism (Figure 1A) to model complex relationships across biological sequences. Pretrained using masked language modeling (MLM), the encoder processes four distinct sequence formats—protein sequences, coding DNA (CDS), and composite NCDS-protein/CDS structures—to capture contextual dependencies between coding regions and their regulatory contexts (Figure 1B, left). During pretraining, 15% of input tokens undergo stochastic masking: 80% replaced with a 〈mask〉 token, 10% substituted with alternative tokens, and 10% retained intact. This strategic corruption forces the model to learn cross-modal relationships, enhancing its capacity to resolve dependencies between protein structural motifs and adjacent non-coding elements while establishing global sequence context. Implementation details are provided in Methods Section A.2.

Post-training analysis reveals the encoder’s ability to decode fundamental biological principles. UMAP visualization of embedding spaces (Fig. S2) demonstrates two critical learned representations: (1) Codon embeddings cluster around their corresponding amino acids, with synonymous codons (e.g., TCT/TCC for serine) forming tight subgroups within amino acid-specific clouds. (2) Amino acid embeddings organize according to physicochemical properties—positively charged residues arginine(R) and lysine(K) occupy adjacent regions distinct from hydrophobic clusters valine(V) and leucine(L), while negatively charged residues aspartate(N) and glutamate(Q) form complementary polarity groups. This hierarchical organization mirrors known biochemical relationships, confirming the model’s capacity to internalize evolutionary constraints governing sequence-structure-function relationships. Notably, the observed spatial alignment between codons and their translated amino acids emerges spontaneously from pretraining without explicit supervision, validating the MLM objective’s effectiveness in capturing biological semantics.

#### 2.1.2 ProDMM Seq2Seq

ProDMM-Seq2Seq advances biological sequence modeling through an integrated encoder-decoder architecture that synergizes bidirectional context understanding with causal generation capabilities. As illustrated in Figure 1B, this framework builds upon the pretrained encoder’s ability to capture protein-DNA interaction patterns while incorporating a task-oriented decoder interconnected via dynamic cross-attention mechanisms. The architecture operates through coordinated processing stages: input sequences—whether protein amino acid chains or partial regulatory elements—are first encoded into latent representations preserving evolutionary constraints, which subsequently guide the decoder’s autoregressive synthesis of biologically coherent outputs.

The model’s pretraining regimen addresses three interconnected biological synthesis challenges fundamental to genetic engineering. Central to its design is the protein-to-coding DNA (CDS) translation task, which converts amino acid sequences into codon sequences. Complementing this core capability, the framework simultaneously generates upstream regulatory elements (pre-NCDS) that maintain promoter architecture requirements—including conserved spacing of transcriptional initiation motifs relative to coding sequences, and generates downstream non-coding regions (post-NCDS) preserving termination signal fidelity and mRNA stability features.

This multimodal training strategy enables ProDMM-Seq2Seq to resolve evolutionary compromises between competing biological requirements. The architecture learns how synonymous codon selection influences transcriptional initiation efficiency through promoter element optimization, while simultaneously adapting protein structural constraints to shape regulatory sequence evolution. Through cross-attention mediated information exchange, the decoder maintains spatial and functional consistency between generated elements—ensuring synthetic promoters align with CDS translation initiation sites, and termination signals coordinate with mRNA secondary structure requirements.

By bridging protein semantics with cis-regulatory logic, ProDMM establishes a new paradigm for precision bioengineering. The framework’s capacity to jointly generate coding sequences and their regulatory contexts enables systematic exploration of design spaces previously constrained by single-modality approaches. This integration proves particularly transformative for metabolic pathway engineering, where coordinated optimization of enzymatic coding sequences with compatible promoter elements can bypass traditional trial-and-error approaches. These advancements herald a shift from modular biological part design to context-aware pathway-scale engineering. ProDMM’s ability to maintain evolutionary consistency while enabling novel combinatorial exploration demonstrates how machine learning can both mimic and extend nature’s optimization strategies—a critical step toward programmable biological systems with enhanced biotechnological capabilities.

### 2.2 Benchmark Results of the Encoder

The ProDMM encoder’s capabilities were rigorously evaluated through a comprehensive benchmarking framework encompassing both unsupervised and supervised biological prediction tasks. As detailed in Figure 1C (middle panel), the model demonstrated robust performance across zero-shot prediction challenges involving DNA sequence properties and protein functional characteristics. Simultaneously, supervised learning evaluations (Figure 1C, left panel) revealed its effectiveness in deriving meaningful representations for downstream predictive applications. Detailed methodologies can be found in the Methods Section A.4 and A.5.

#### 2.2.1 Zero-Shot Tasks

The ProDMM encoder’s architectural design enables systematic resolution of biological sequence interdependencies, capturing intrinsic semantic relationships between coding DNA (CDS), protein sequences, and flanking non-coding regulatory elements (NCDS). This capability extends to modeling inter-molecular interactions critical for deciphering genetic regulatory networks, particularly the co-evolutionary dynamics between promoter regions (upstream NCDS) and their associated protein-coding sequences.

We hypothesized that the encoder’s integrated representation space would support accurate zero-shot prediction of: (1) sequence-specific physicochemical properties, (2) cross-modal functional interactions, and (3) pathway-level regulatory outcomes. To validate this hypothesis, a comprehensive benchmarking protocol was implemented comparing ProDMM against state-of-the-art protein- and DNA-specific models across six biological prediction tasks.

##### Zero-shot CDS expression prediction

We present that the perplexity of CDS sequences calculated by the ProDMM-Encoder (for the method of calculating sequence perplexity, see Method Section A.4 and A.4.1) correlates with the expression levels of CDS sequences. To validate this, We constructed a dataset from literature [11], which contains expression levels from five different proteins. Each dataset records the expressive levels measured in experiments for different codons of the same protein. Figure 2A displays the Spearman rank correlation between the CDS perplexity and the experimental expression level, while each dot corresponds to a specific dataset. The results show a strong correlation between the perplexity and expression level (Average Spearman *r* = 0.8, two-sided t-distributed *p <* 1 × 10^−5^), indicating that lower perplexity corresponds to higher yield. We also compare ProDMM with alternative nucleotide baseline models, including Evo [7], RNA-FM [12], and GenSLM 2.5B [13]. ProDMM exhibits the best performance, demonstrating its superior ability to model gene expression accurately.

**Figure 2:**
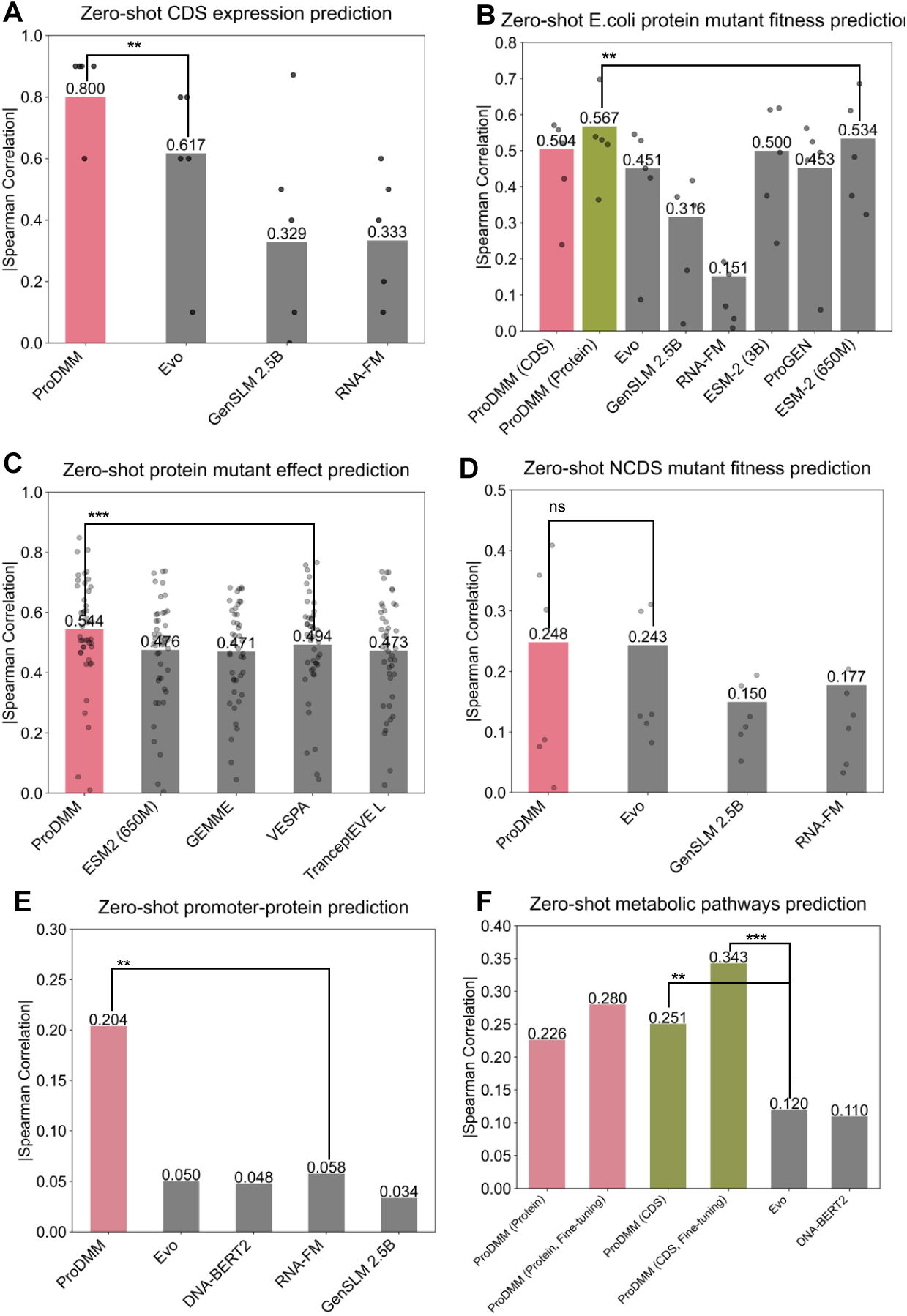
ProDMM performs zero-shot function prediction for CDS, proteins, NCDS, promoter-protein, and metabolic pathways. (A-D) Zero-shot prediction performance of ProDMM-Encoder. The bar height represents the Spearman rank correlation between the target sequence perplexity calculated by ProDMM-Encoder and the experimental fitness (expression level, activity, stability, etc.), while each dot corresponds to a specific dataset. (E) The Spearman correlation between the perplexity of promoter-protein complex sequence and expression level. (F) The Spearman correlation between the perplexity of metabolic (NCDS-Protein-NCDS-Protein-NCDS-Protein-NCDS-Protein) complex sequence and the experimental expression level. P-values (two-side t-test) are reported between the top two performing models; The asterics indicate the p-value, ranging from 0.01-0.05 (*), 0.001-0.01 (**), 0.0001-0.001 (***).

##### Zero-shot E.coli protein mutant fitness prediction

ProDMM-Encoder can predict the impact of CDS mutations on protein fitness. Specifically, the scoring of the mutant by ProDMM-Encoder is correlated with the fitness of the mutants (for the method of calculating the mutant score, refer to Method Section A.4 and A.4.3). We used a deep mutation scanning (DMS) dataset containing mutations from five proteins from E. coli, which was compiled by Evo [7]. Since the input of ProDMM-Encoder can be either protein or CDS, we separately tested the performance using protein as input and using CDS as input. As shown in Figure 2B, when ProDMM utilizes protein sequences as input, it demonstrates superior performance compared to other protein sequence models, such as ESM2 and progen [5, 14]. Conversely, when using CDS DNA sequences, ProDMM similarly excels against other nucleotide sequence models, including Evo [7], GenSLM 2.5B [13] and RNA-FM [12]. Moreover, ProDMM’s performance with protein sequence input consistently surpasses that with CDS sequence input in this task.

##### Zero-shot protein mutant fitness prediction

ProDMM-Encoder, pre-trained with masked language modeling, can serve as a protein mutant fitness predictor [15] (for the method of calculating the mutant score, refer to Method Section A.4 and A.4.4). Given that ProDMM’s training set is derived from the GTDB [16] and only comprises prokaryotic genomes, Figure 2C shows the ProDMM’s performance in the mutation-fitness prediction task against all 45 prokaryotic protein datasets within the ProteinGym benchmark [17]. ProDMM consistently outperformed other baseline sequence models, including ESM2 [5], GEMME [18], VESPA [19] and TranceptionEVE [20], affirming its robustness and reliability in predicting evolutionary fitness under various genetic contexts.

##### Zero-shot NCDS mutant fitness prediction

In Figure 2D, we assess the zero-shot prediction capabilities of ProDMM concerning non-coding RNA (ncRNA), leveraging data obtained from Evo’s research [7]. In this endeavor, ProDMM employs its NCDS modality to input DNA sequences (translated from ncRNA) and once again outperforms other nucleotide models, including GenSLM [13] and RNA-FM [12], thereby reinforcing the model’s comprehensive predictive abilities.

##### Zero-shot promoter-protein prediction

Moreover, in Figure 2E, we conduct an in-depth examination of a zero-shot prediction task targeting promoter transcription intensity in E. coli, drawing upon data found in the literature [21]. In this study, the authors investigated the fluorescence intensity of EGFP under diverse promoter conditions, thereby generating insights into transcriptional efficiencies. We provided ProDMM with both the NCDS and the corresponding protein sequence. Interestingly, although the EGFP protein sequence remained constant throughout the experimental evaluations, ProDMM’s predictions of transcriptional intensity significantly outperformed those of other nucleotide baseline models. This finding suggests that ProDMM has adeptly captured the complex interactions between promoter sequences and protein sequences, highlighting their critical role in accurately predicting the transcriptional efficiency of downstream proteins—an aspect that has often been insufficiently explored in previous studies.

##### Zero-shot metabolic pathways prediction

Building upon the above findings, we propose that ProDMM, having assimilated an extensive array of interaction features between proteins and their upstream and downstream non-coding sequences during its pre-training phase, is endowed with the capacity for zero-shot predictive capabilities regarding metabolite yields in metabolic pathways. To validate this proposition, we leveraged data from the existing literature [4], wherein authors constructed a sophisticated metabolic pathway in E. coli consisting of four distinct proteins and four strategically selected promoter sequences designed to facilitate the biosynthesis of naringenin from L-tyrosine. The authors employed a suite of experimental methodologies to quantitatively assess the yield of naringenin produced by these constructed pathways, investigating various promoter combinations while maintaining constant protein sequences.

To thoroughly evaluate the performance of ProDMM in this metabolic context, we employed the Perplexity-based Zero-shot Sequences Ranking method using the ProDMM-encoder, as delineated in the methods section. This approach permitted the direct scoring of different pathway sequences comprising eight components formed by the four NCDS coupled with their corresponding protein sequences. We subsequently assessed the scores in relation to the yield of the final product, naringenin. As depicted in Figure 2F, the zero-shot predictive performance of the pre-trained ProDMM outperforms alternative nucleotide models (two-sided t-distributed *p <* 0.01), including Evo and DNABERT2. This remarkable distinction underscores the unique capabilities of ProDMM.

During its pre-training phase, ProDMM acquired knowledge regarding the interactions among individual protein sequences (or coding sequences, CDS) alongside their associated upstream and downstream non-coding sequences. However, this phase did not directly address long-range interactions among various protein components that are distributed throughout metabolic pathways. In contrast, other nucleic acid models such as Evo [7] and DNABERT2 [8] have leveraged information from the entire genome, thereby capturing longer-range interactions among CDS sequences to a greater extent. To further enhance the model’s predictive capabilities further, we meticulously compiled and organized genomic and protein data from the antiSMASH database version 4 [22], encompassing 36,554 species. This dataset was structured into sequences formatted as NCDS-protein-NCDS-protein, containing four components per sequence. Following this, we fine-tuned ProDMM’s encoder on this structured data using a masked language model (MLM) pretraining approach, enabling ProDMM to learn the interactions between adjacent proteins and their corresponding non-coding sequences. When we assessed the zero-shot predictive performance of the fine-tuned model on the aforementioned metabolic pathway dataset, we observed a significant enhancement in predictive accuracy. This finding underscores the effectiveness of explicitly training ProDMM to comprehend the interactions between neighboring proteins and their non-coding sequences, thereby amplifying its zero-shot predictive capabilities for more complex and extended metabolic pathways.

We anticipate that extending ProDMM’s architecture to learn longer-range interactions between protein sequences will further improve its zero-shot predictive abilities concerning metabolic pathways. By providing invaluable insights into the complexities associated with metabolic pathway optimization, ProDMM establishes a robust framework for future research endeavors in this critical area. Ultimately, the work presented here not only advances our understanding of genetic interactions but also opens new avenues for innovative biotechnological applications and sustainable bioproduction processes. This contribution significantly enhances the broader fields of synthetic biology and metabolic engineering, promoting ongoing exploration and growth in these transformative disciplines.

#### 2.2.2 Supervised Learning Tasks

We further evaluated the performance of ProDMM-encoder across a diverse array of supervised learning tasks, specifically focusing on its ability to predict melting temperatures for various native proteins (Figure 3A), forecast protein solubility (Figure 3B), estimate protein expression levels (Figure 3C), and assess the transcriptional abundance of proteins in Escherichia coli and Halobacterium volcanii (Figure 3D). To ensure consistency with prior studies, we utilized the Pearson correlation coefficient as the evaluation metric for these supervised learning tasks [23]. In each of these tasks, ProDMM consistently demonstrated superior performance when utilizing coding sequences (CDS) as input compared to utilizing protein sequences. Furthermore, for most of the tasks, ProDMM outperformed several established protein and nucleotide baseline models, including CaLM [23], ESM [5, 15, 24], and ProtTrans [6], underscoring its efficacy in handling these challenges. Comprehensive data can be found in Table S1 of the supplementary information.

**Figure 3:**
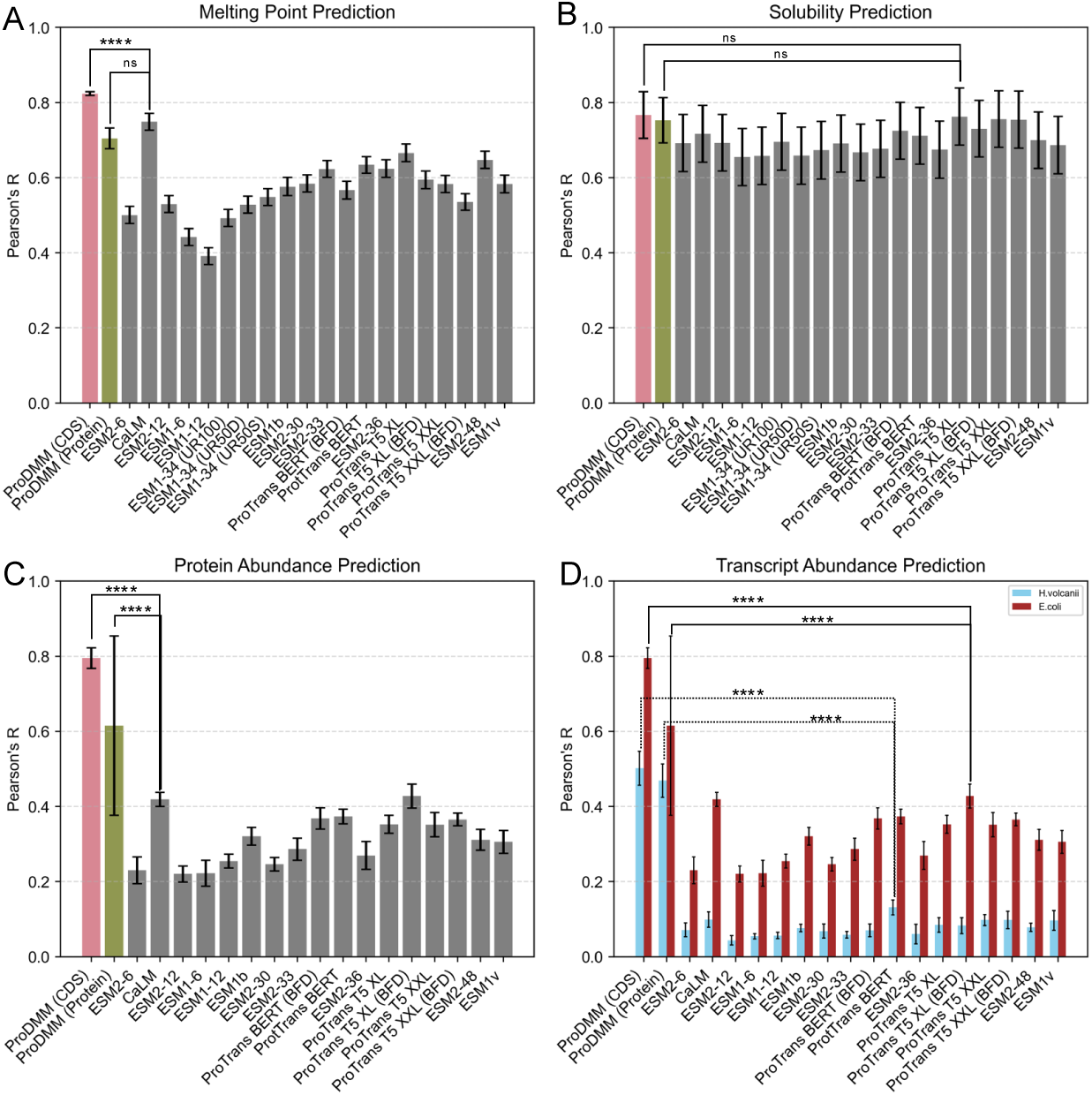
Evaluation of ProDMM-Encoder in protein property prediction tasks. This figure assesses the performance of the ProDMM model across four protein property prediction tasks. (A)-(D) show the results for melting point prediction, solubility prediction, protein abundance prediction, and transcript abundance prediction, respectively. P-values are reported between the top three performing models; “****” denotes *p <* 0.0001. Performance is measured using the Pearson correlation coefficient, reflecting the correlation between predicted and actual values. The data are presented as mean values, with error bars representing the standard deviation calculated from a fivefold cross-validation. The results from the ESM and ProtTrans series of models were obtained from the literature [23].

Additionally, we visualized the embeddings generated by ProDMM for the top ten organisms with the highest volume of sequence data, utilizing both CDS sequences and protein sequences as input, as depicted in Figure S3. The visual representations derived from these figures provide compelling evidence that embeddings obtained from CDS sequences exhibit sharper and more distinct boundaries across various species compared to those generated from protein sequences. This observation underscores the model’s capability to delineate subtle differences in sequence information when CDS sequences are employed, suggesting that they encapsulate richer and more nuanced biological insights.

Collectively, these findings underscore the remarkable capabilities of the ProDMM model, particularly when leveraging CDS sequences. This enhanced performance across the aforementioned protein-related supervised learning tasks indicates that ProDMM is adept at capturing a comprehensive spectrum of biological information inherent in the sequences, thereby improving predictive accuracy and reliability. The ability to integrate such diverse data effectively may pave the way for further advancements in protein research and synthetic biology applications, providing a robust foundation for future explorations in the field. By harnessing the intricate details within CDS sequences, ProDMM not only elevates the quality of predictions but also contributes significantly to our understanding of protein behavior and function across different biological contexts.

### 2.3 Benchmark Results of the Generation

To evaluate the performance of the ProDMM Seq2Seq, we built up and conducted a series of benchmarks across multiple downstream tasks, including prokaryotic and eukaryotic organisms reverse translation of proteins, as well as various promoter-cds co-generation tasks (Figure 4). Detailed methodologies can be found in the Methods section.

**Figure 4:**
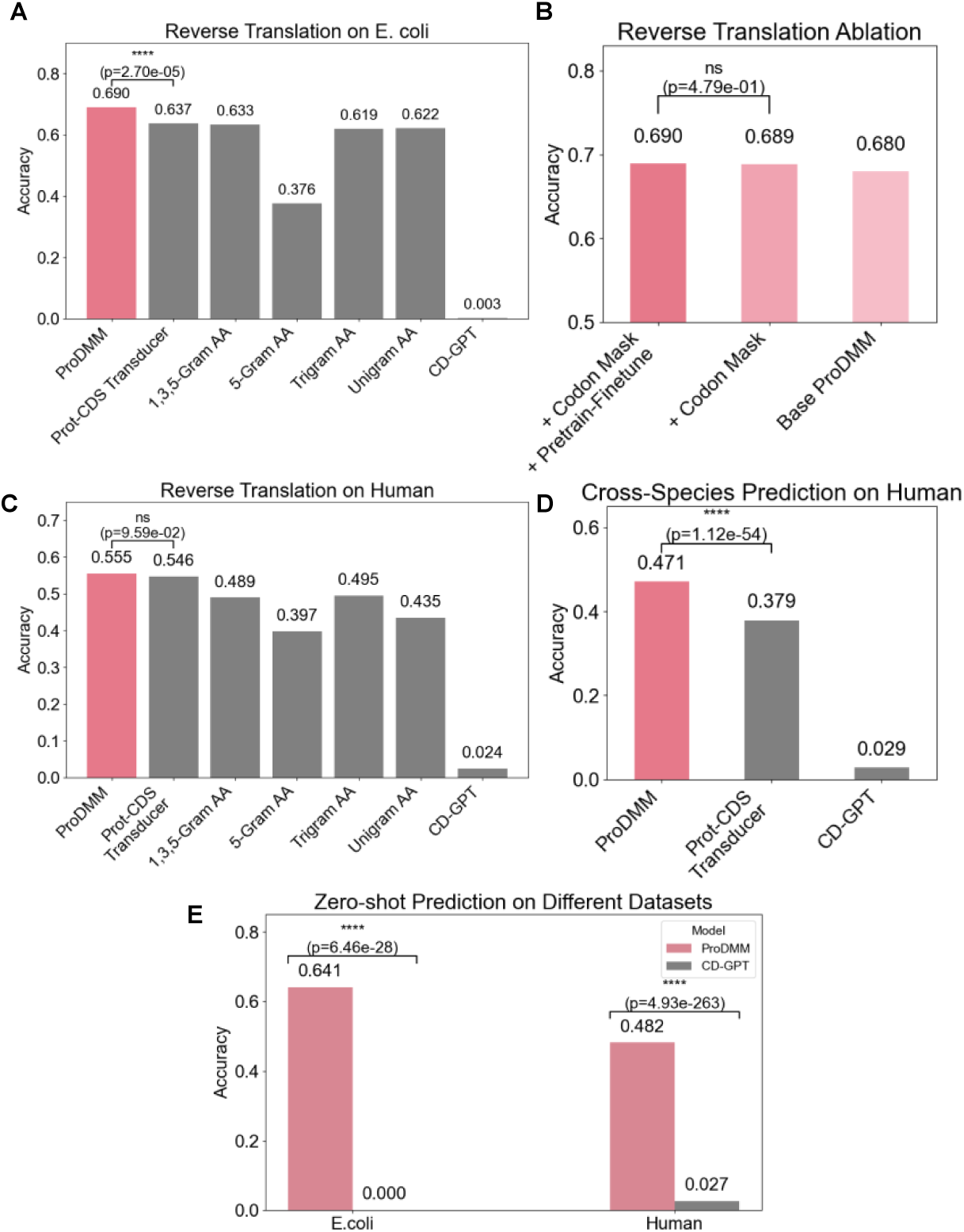
Evaluation of ProDMM on Reverse Translation Tasks. This figure assesses the performance of the ProDMM model across various reverse translation tasks. (A) displays the results for the reverse translation of high-expression CDS based on a protein sequence from E. coli. (B) shows the impact of ablation studies on codon mask constraints and the pretrain-finetune approach for reverse translation tasks. (C) illustrates the reverse translation of high-expression CDS based on a protein sequence from humans. (D) demonstrates cross-species prediction, where the model is fine-tuned on the E. coli dataset and tested on the human dataset. (E) evaluates the zero-shot prediction capabilities of reverse translation without any fine-tuning. Performance is measured using accuracy, reflecting the correlation between the generated codon sequences and the actual codon sequences.

#### 2.3.1 Reverse Translation Generation

ProDMM’s sequence-to-sequence (seq2seq) architecture intricately investigates the inherent relationships among proteins, CDS, and NCDS, thereby enhancing its cross-modal generative capabilities. In Figure 4, we validate the efficacy of this cross-modal generation by focusing on the task of generating codon sequences that correspond accurately to given protein sequences. This involves the reverse translation of proteins while implementing codon mask constraints. These constraints are crucial, as they ensure that the generated codons precisely match the specific amino acids, adhering to the standard codon table. This methodological approach highlights the model’s proficiency in executing complex sequence transformations while maintaining strict biological fidelity.

We adopt the dataset from Transducer [25], where the task is to generate high-expression CDS given a protein sequence. As shown in Figure 4A, on the high-expression data from E. coli, ProDMM outperforms baseline models based on unigram, trigram, and 5-gram frequencies, as well as their ensemble (1,3,5-gram) [26], and significantly surpasses the state-of-the-art Prot-CDS Transducer model. Additionally, Figure S4 demonstrates that ProDMM accurately produces the correct codon usage distribution for E. coli. In Figure 4B, we conducted ablation studies on the E. coli data, including the incorporation of codon mask constraints and a pretrain-finetune approach. Due to the scarcity of high-expression gene data in E. coli, we hypothesized that pretraining on a larger set of non-high-expression gene data from E. coli, followed by finetuning on high-expression genes, would be beneficial—forming a pretrain-finetune pipeline. Experimental results confirmed that adding amino acid-codon mask constraints and employing the pretrain-finetune strategy both contribute positively to the generation performance.

Interestingly, as shown in Figure 4C, on high-expression human data, ProDMM’s generated results also outperform those of the baseline models. Despite being pre-trained on a large amount of prokaryotic data, ProDMM can effectively transfer to eukaryotic data without experiencing model collapse, demonstrating that extensive pre-training has endowed the model with species adaptability. Furthermore, we conducted experiments without finetuning on human data to explore the capability of cross-species high-expression generation. As shown in Figure 4D, we directly generalized ProDMM and Prot-CDS Transducer—both finetuned on high-expression E. coli data—to the generation of high-expression human CDS. We found that ProDMM’s cross-species generalization ability significantly surpasses that of Transducer. Similarly, as shown in Figure 4E, when we do not perform any finetuning on high-expression data and use only the pre-trained model for zero-shot prediction, ProDMM already outperforms the baseline biological model CD-GPT [27] on both E. coli and human data. This reflects the effectiveness of large-scale cross-modal pre-training in understanding and generating the intrinsic relationships between proteins and CDS.

#### 2.3.2 Regulator-CDS Co-evolving Effect

Some studies [28, 29] have indicated that there exists a co-evolving interaction between gene regulatory elements and CDS, jointly influencing gene expression. Using the E. coli data from DeepExpression [29], we trained a model called ProDMM-Expression. This model utilizes the ProDMM encoder to predict gene expression from DNA sequences, encompassing both coding and non-coding regions. In our approach, we first conditioned on the CDS to generate specific promoters, which were then concatenated with the CDS and fed into ProDMM-Expression. For control experiments, we separately used only the Ground Truth (GT) CDS or only the promoter as input. As depicted in Figure 5A, the tailored promoters, specifically designed to complement the CDS, when combined with the CDS, significantly enhance gene expression prediction. This method achieves superior results compared to using the CDS or the promoter alone, underscoring the synergistic effect of customized promoter-CDS combinations in predicting gene expression.

**Figure 5:**
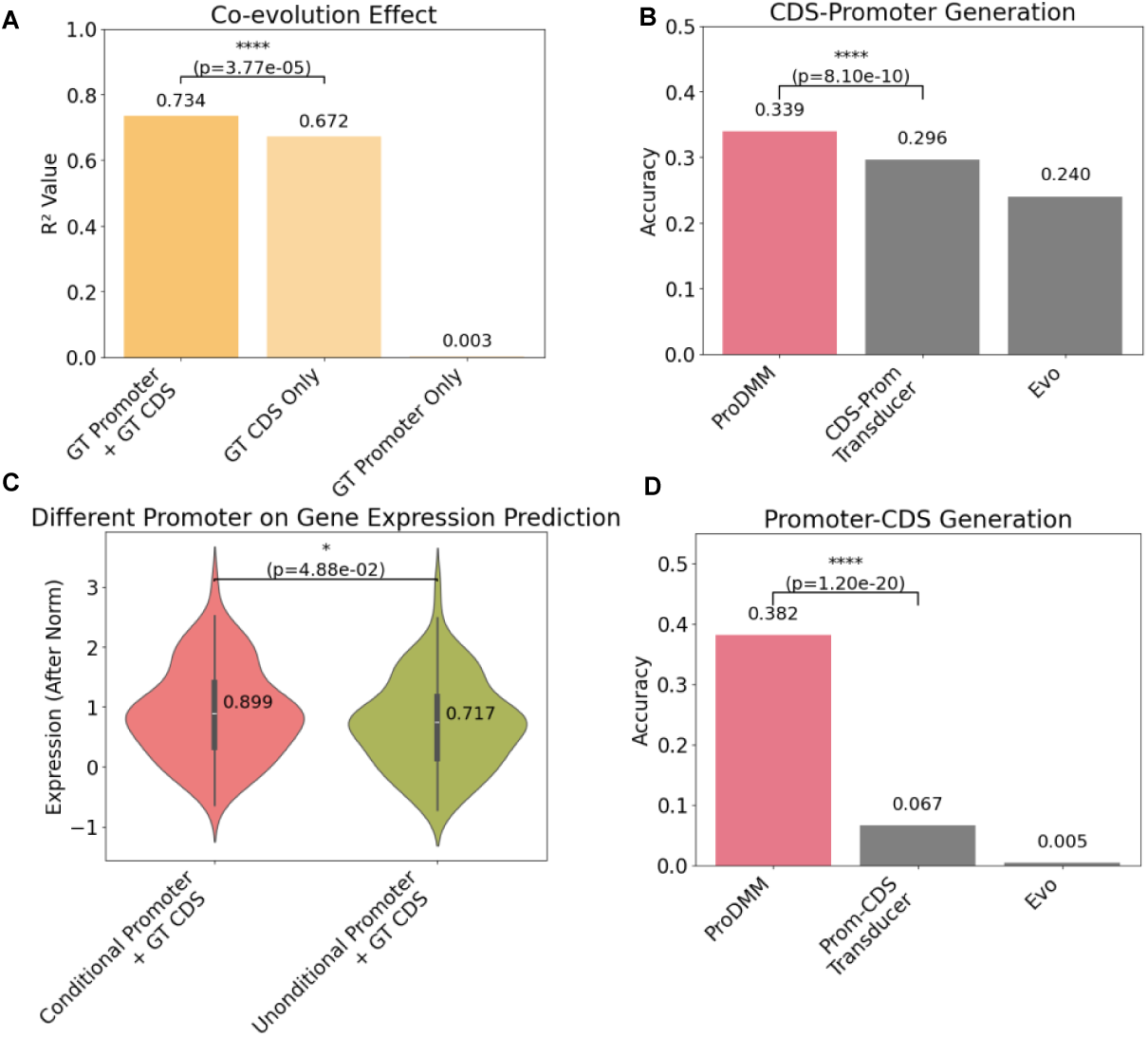
Evaluation of ProDMM on Co-evolution Effect Tasks. This figure assesses the performance of the ProDMM model across reverse translation tasks. (A)-(D) show the results for the co-evolution effect of the CDS and the promoter, CDS to promoter generation, comparison between conditional and unconditional promoter for gene expression, and promoter to CDS generation, respectively.

Given the significant impact of promoter specificity on protein expression [30], we have conducted an in-depth investigation into generating optimal promoters based on CDS information. To achieve this, we curated a dataset of promoter-CDS pairs of E. coli for model fine-tuning [31]. As illustrated in Figure 5B, ProDMM significantly outperforms existing methods, including the DNA-based approach Evo [7] and the baseline method CDS-Prom Transducer. Despite the availability of numerous methods for unconditional promoter generation, there is a conspicuous gap in research focusing on generating specific promoters conditioned on CDS. To emphasize the importance of conditioned promoter generation, we conducted a series of comparative experiments. These studies juxtaposed conditionally generated promoters combined with CDS sequences against unconditionally generated ones, leveraging ProDMM-Expression as a reference for expression level predictions (Figure 5C). The findings indicate that promoters developed with CDS-specific conditioning yield substantially higher gene expression levels relative to those generated unconditionally. Additionally, Figure S5 illustrates that conditional generation successfully reproduces the distribution of key motifs in promoter sequences, such as the CCAAT box, the -75 region, and the TATAAT box in the -10 region, more effectively than unconditional generation. This not only accentuates the essential need to tailor promoter generation to CDS for improved gene expression but also showcases the pivotal role of ProDMM’s advanced cross-modal generative capabilities in propelling this research frontier forward.

Building upon our previous co-evolution analysis, which established the specific and intricate relationship between CDS and promoters, we have embarked on an exploration of a novel task: the generation of CDS sequences guided by promoters. The objective is to produce CDS that more effectively harness the functional capabilities of their associated promoters. In this endeavor, we compared our model, ProDMM, with baseline models Evo and Prom-CDS Transducer. As illustrated in Figure 5D, ProDMM’s proficiency in generating promoter-specific CDS significantly surpasses that of the other two baseline approaches. Notably, unlike the reverse translation and CDS-guided promoter generation tasks, the Promoter-Specific CDS Generation task was not part of the model’s pre-training regimen. This emphasizes the robustness of ProDMM’s cross-modal generation capabilities, which facilitate the exploration of interactions across different modalities. Furthermore, this highlights ProDMM’s ability to expand the frontiers of cross-modal generation, showcasing its potential to innovate in the domain of gene regulatory mechanisms.

## 3 Discussions

In this study, we present ProDMM, a cutting-edge multimodal protein-DNA model pre-trained on over 129 million pairs of non-coding DNA sequences (NCDS) and coding sequences (CDS). This innovative architecture supports three distinct input modalities in one entry—protein sequences, CDS, and NCDS—allowing for versatile applications across various biological contexts. At its core, ProDMM consists of a BERT-like encoder complemented by a decoder tailored for numerous generative tasks.

ProDMM has demonstrated extraordinary performance across a diverse array of downstream tasks, including zero-shot predictions for protein sequences, CDS sequences, NCDS sequences, and CDS expression levels. It excels in supervised learning tasks, accurately predicting the melting temperature *T_m_* and solubility of native protein sequences. Additionally, it performs exceptionally well in conditional generation tasks—generating CDS sequences based on protein sequences and promoter sequences based on CDS—far surpassing existing baseline models in both protein and nucleic acid prediction, such as ESM [5, 15, 24] and ProtTrans [6] for proteins, and Evo [7], DNABERT2 [8] for nucleic acids. Furthermore, we discovered that in supervised learning tasks related to protein sequences, utilizing the CDS as input provided superior predictive performance compared to using amino acid sequences. In promoter generation tasks, conditional generation based on CDS sequences yielded substantially higher protein expression levels compared to unconditional approaches. These findings underscore the significant implications for future initiatives in protein engineering and promoter design, highlighting the necessity for focused strategies in these domains. A key strength of ProDMM lies in its architectural design, which effectively captures the intricate interactions between protein sequences and their upstream and downstream NCDS. Notably, ProDMM can predict the characteristics of entire metabolic pathways—specifically, the yield of end products—without any reliance on labeled training data, thus showcasing its impressive zero-shot predictive capabilities. Moreover, supplementary training on data from adjacent protein sequences and their associated NCDS substantially enhances ProDMM’s zero-shot predictive performance concerning metabolic pathways. In conclusion, ProDMM represents a transformative advance in the fields of synthetic biology, protein engineering, promoter design, codon optimization, and metabolic pathway optimization. Its versatility and predictive power open avenues for innovative strategies and improved outcomes in metabolic engineering and biotechnological applications, heralding a new era of advancements in these vital scientific disciplines.

## Ethics approval and consent to participate

Not applicable.

## Consent for publication

Not applicable.

## Author contribution

P.T., L.H. and N.D. conceptualized and supervised this research project. P.T., M.L., Y.R. and P.Y. developed the architecture of ProDMM. P.T. collected and processed the pre-training dataset. M.L., Y.R., P.Y. and J.C. wrote the code and conducted the pre-trianing of the model. P.T. designed the benchmarks. M.L., Y.R., P.Y., J.C., N.D., X.M., Y.C., B.Z., W.O. and B.W. finished the benchmarks. M.L., Y.R., P.Y. and J.C. contributed equally to this work. All authors reviewed and accepted the manuscript.

## Appendix A Method

### A.1 Pre-training Dataset Processing

The pre-training dataset was sourced from the GTDB database, release number 220. To reduce the data volume, we only utilized sequences from representative genomes. We then employed the Prodigal [32] tool to predict the CDS sequences from each nucleotide FASTA file and obtained the corresponding translated protein sequences. During this process, we also recorded the non-coding sequences (NCDS) located between adjacent CDS sequences within the same FASTA file. This approach allowed us to retrieve each CDS sequence along with its preceding and post-NCDS sequences, as well as the translated protein sequences. During the processing, we discarded any non-full-length protein sequences and protein sequences exceeding 2000 amino acids in length. Additionally, protein sequences that were not translated using the standard codon table or contained non-standard amino acids were excluded. Following these filtering steps, we ultimately retained approximately 129 million paired sequences.

### A.2 ProDMM-Encoder

ProDMM-Encoder includes a universal tokenizer that converts protein sequences, coding sequences (CDS), and non-coding sequences (NCDS) into one-hot encodings. It features a Transformer encoder that is pre-trained using a masked language modeling (MLM) task to derive sequence latent representations. Additionally, it has a prediction head that generates token distributions from these latent representations, enabling ProDMM-Encoder to perform zero-shot sequence scoring tasks.

#### Universal tokenizer

The universal tokenizer splits a molecule sequence (Protein, CDS, or NCDS) into a small tokens list. It then maps each token to a unique integer index using a vocabulary table, followed by transforming these indices into one-hot vectors with all elements set to 0 except for a 1 at the position corresponding to the index of the token. In protein sequences and NCDS, each character is a token while every three characters form a token in the CDS. The vocabulary consists of 97 tokens: 64 codons, 20 natural amino acids, 4 nucleotide bases (A, T, C, G), and 9 special tokens. To distinguish between the abbreviations of amino acids and nucleotide bases in the program implementation, we use uppercase characters to represent amino acids and lowercase characters to represent nucleotide bases. For instance, uppercase ‘**A**’ represents *alanine*, while lowercase ‘**a**’ represents *adenine*; uppercase ‘**T**’ stands for *threonine*, and lowercase ‘**t**’ denotes *thymine*. The tokens, indices, and names in the vocabulary are listed in Table A1.

#### Model Architecture

The ProDMM is based on the Transformer architecture [9], but we have made several modifications that have been proven effective in practice. Specifically, we replaced absolute position encoding with rotary position encoding [33], replaced the ReLU activation function with the GeLU activation function [34], replaced Pre-LayerNorm with Post-LayerNorm [35], optimized the standard attention operation using flash attention [36], added a layer normalization layer after the embedding [10], and used embedding shrinking [37] technique to prevent loss spike.

Formally, given a molecular sequence, it is tokenized into a sequence of tokens (*t*_1_*, t*_2_*, . . ., t_N_* ), where *N* denotes the sequence length. Each token *t*_2_ is mapped to a unique integer index *x_i_* according to the vocabulary. These indices are then transformed into one-hot vectors **e***_i_* ∈ ℝ*^V^*, where *V* is the size of the vocabulary. In each one-hot vector **e***_i_*, all elements are zero except for a single one at the position corresponding to the index *x_i_*.

The one-hot vectors are projected into a continuous embedding space using an embedding matrix **E** ∈ ℝ*^V^* ^×*d*^, where *d* is the dimensionality of the embedding space:

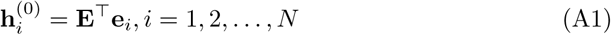

These embeddings are fed into a stack of *L* Transformer encoder layers. At each layer *l* 1 ≤ *l* ≤ *L*), the hidden representations are updated using multi-head self-attention mechanisms followed by position-wise feed-forward networks. The computation within each Transformer layer can be summarized as:

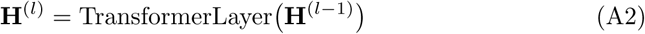

where 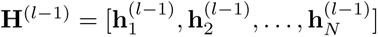 represents the matrix of hidden states from the previous layer.

After passing through all *L* layers, we obtain the final contextualized representations **H**^(^*^L^*^)^. For the masked language modeling task, these representations are projected back to the vocabulary space to compute the logits for each token position:

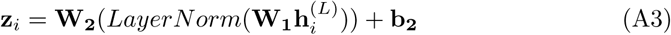

where **W_1_**∈ ℝ*^d^*^×*d*^ is a projection matrix, **W_2_**∈ ℝ*^V^* ^×*d*^ is the output projection matrix and **b_2_** ∈ ℝ*^V^* is a bias vector.

The predicted probability distribution over the vocabulary at each position is obtained by applying the softmax function to the logits:

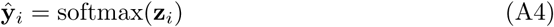

**Table A1:**
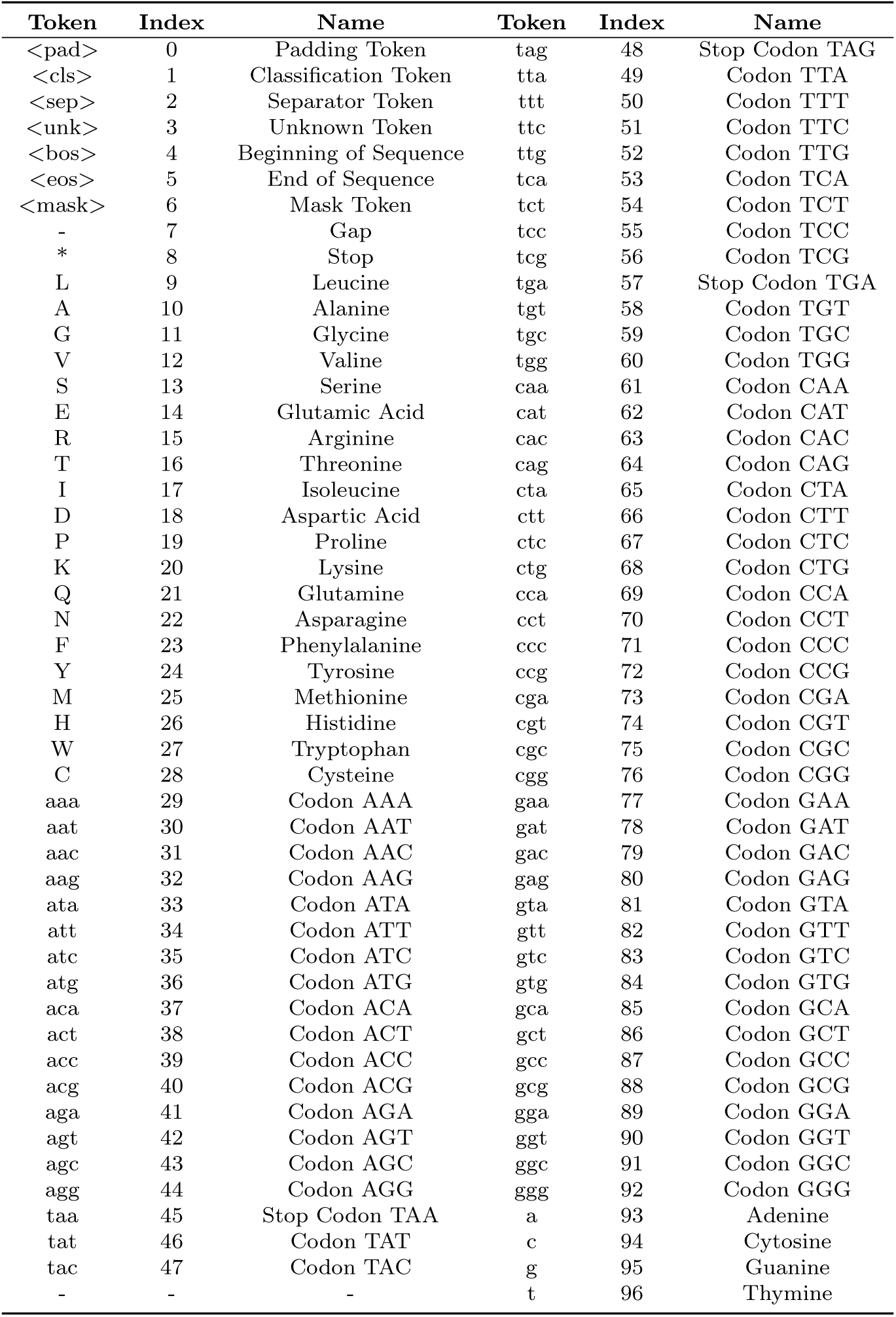
Vocabulary table of tokens, indices, and names.

The model is trained by minimizing the cross-entropy loss between the predicted distributions and the true one-hot encoded targets **y***_i_*:

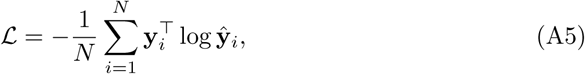

#### Pre-training Method

ProDMM-Encoder is pre-trained with masked language modeling (MLM) on the following sequences:

- Protein Sequence. *<*cls*>* token and *<*sep*>* token are added at the beginning and end of tokenized sequences.
- CDS Sequence. *<*cls*>* token and *<*sep*>* token are also added at the beginning and end of tokenized sequences.
- NCDS-CDS-NCDS Sequence. *<*cls*>* token and *<*sep*>* token are added at the beginning and end of tokenized sequences. Additionally, a *<*sep*>* token is inserted between NCDS and CDS, i.e., “{*<*cls*>*, NCDS tokens, *<*sep*>*, CDS tokens, *<*sep*>*, NCDS tokens, *<*sep*>*}”.
- NCDS-Protein-NCDS Sequence. *<*cls*>* token and *<*sep*>* token are added at the beginning and end of tokenized sequences. Additionally, a *<*sep*>* token is inserted between NCDS and Protein, i.e., “{*<*cls*>*, NCDS tokens, *<*sep*>*, Protein tokens, *<*sep*>*, NCDS tokens, *<*sep*>*}”

Each *<*pre-NCDS, CDS, Protein, post-NCDS*>* quadruple in the dataset transitions to one of the sequences above with equal probability.

For each sequence, 15% of the tokens are randomly selected, of which 80% are replaced with the mask token, 10% are randomly replaced with a random token, and 10% are left unchanged [10]. The model is trained using cross-entropy loss in Equation A4. We opted to use the 15% masking ratio because this ratio has been extensively validated as the most effective in pre-training models for proteins [38], DNA [8, 23, 39], and molecules [40]. We utilized AdamW optimizer [41, 42] to train the model with a learning rate of 1.5 × 10^−4^, *β*_1_ = 0.9, *β*_2_ = 0.95, *ɛ* = 10^−8^, and a weight decay of 0.01. We warmed up the learning rate gradually during the initial 5,000 gradient update steps, starting from 0 and reaching 1.5 × 10^−4^. Subsequently, we apply a cosine learning rate scheduler to decay the learning rate over the remaining steps. To monitor the training process, 5% of the training set was randomly selected for validation purposes. We trained the model with 24 NVIDIA H100 GPUs with 80GB VRAM memory over 40 days, which involved 260,000 gradient steps. We manually terminated the training as there was no improvement in validation loss for 10,000 training steps. We employed the zero-1 strategy within the DeepSpeed framework [43] to distribute optimizer parameters across GPUs for enhanced computational efficiency. We did not zero-2 and zero-3 due to the additional communication overhead they introduce when gradient accumulation is in use. The training curves for both the encoder training loss and validation loss are presented in Figure S6. The hyper-parameters configuration are listed at Table A2.

**Table A2:** Pre-training hyper-parameters of ProDMM-Encoder.

| Parameter | Value |
| --- | --- |
| Dropout | 0.1 |
| Hidden size | 1536 |
| Intermediate size | 6144 |
| LayerNorm epsilon | $1.0 \times 10^{-5}$ |
| Number of attention heads | 64 |
| Number of hidden layers | 48 |
| Training mini-batch size | 4096 |
| Evaluation mini-batch size | 4096 |
| Seed | 42 |
| Training steps | 260,000 |
| Gradient Accumulations | 8 |
| Learning rate | $1.5 \times 10^{-4}$ |
| Adam beta1 | 0.9 |
| Adam beta2 | 0.95 |
| Adam epsilon | 1e-8 |
| Gradient clip value | 1.0 |
| Gradient clip algorithm | norm |
| Precision | BF16 |
| Weight decay | 0.01 |
| Warmup steps | 5000 |
| Strategy | Zero-1 |

### A.3 ProDMM-Seq2Seq

ProDMM-Seq2Seq includes a prompt tokenizer based on the universal tokenizer in ProDMM-Encoder that expands cross-modality generation prompt tokens. It enables sequence-to-sequence (seq2seq) generation using a language modeling (LM) task to encode source sequence latent representations, then decode it to target sequence by using a casual decoder.

#### Prompt tokenizer

We expanded the universal vocabulary which originally contains 97 tokens, by adding three generation prompt tokens. These prompt tokens are designed to guide the model to perform cross-modal generation tasks more effectively. The vocabulary consists of 100 tokens: 9 special tokens, 64 codons, 20 natural amino acids, 4 nucleotide bases (A, T, C, G), and 3 generation prompt tokens.

#### Pre-training Method

ProDMM-Seq2Seq is pre-trained with language modeling (LM) on the following sequences:

- Protein Sequence - CDS. **Source Sequence:** Protein Sequence - *<*cls*>* token and *<*sep*>* token are added at the beginning and end of tokenized sequences. Additionally, a *<*aas2cds*>* token is inserted between Protein Sequence and *<*sep*>*, i.e., “{*<*cls*>*, Protein tokens, *<*aas2cds*>*, *<*sep*>*}”. **Target Sequence:** CDS - *<*bos*>* token and *<*eos*>* token are added at the beginning and end of tokenized sequences, i.e., “{*<*bos*>*, CDS tokens, *<*eos*>*}”.
- CDS - Pre-NCDS. **Source Sequence:** CDS - *<*cls*>* token and *<*sep*>* token are added at the beginning and end of tokenized sequences. Additionally, a *<*cds2pre-ncds*>* token is inserted between Protein Sequence and *<*sep*>*, i.e., “{*<*cls*>*, CDS tokens, *<*cds2pre-ncds*>*, *<*sep*>*}”. **Target Sequence:** Pre-NCDS - *<*bos*>* token and *<*eos*>* token are added at the beginning and end of tokenized sequences, i.e., “{*<*bos*>*, Pre-NCDS tokens, *<*eos*>*}”.
- CDS - Post-NCDS. **Source Sequence:** CDS - *<*cls*>* token and *<*sep*>* token are added at the beginning and end of tokenized sequences. Additionally, a *<*cds2post-ncds*>* token is inserted between Protein Sequence and *<*sep*>*, i.e., “{*<*cls*>*, CDS tokens, *<*cds2post-ncds*>*, *<*sep*>*}”. **Target Sequence:** Post-NCDS - *<*bos*>* token and *<*eos*>* token are added at the beginning and end of tokenized sequences, i.e., “{*<*bos*>*, Post-NCDS tokens, *<*eos*>*}”.

**Table A3:** Seq2Seq prompt vocabulary table of tokens, indices, and names based on the universal vocabulary of the encoder.

| Token | Index | Name |
| --- | --- | --- |
| <aas2cds> | 97 | Amino Acids Sequence to CDS Token |
| <cds2pre-ncds> | 98 | CDS to Pre-NCDS Token |
| <cds2post-ncds> | 99 | CDS to Post-NCDS Token |

We utilized AdamW optimizer [41, 42] to train the model with a learning rate of 4 × 10^−4^, *β*_1_ = 0.9, *β*_2_ = 0.999, *ɛ* = 10^−7^, and a weight decay of 0.01. We warmed-up the learning rate gradually during the initial 5,000 gradient update steps, starting from 0 and reaching 1.5×10^−4^. Subsequently, we apply a cosine learning rate scheduler to decay the learning rate over the remaining steps. To monitor the training process, 5% of the training set was randomly selected for validation purposes. We trained the model with 32 NVIDIA A100 GPUs with 80GB VRAM memory over 14 days, which involved 100,000 gradient steps. We manually terminate the training as there was no improvement in validation loss for 5,000 training steps. We employed DDP strategy for enhanced computational efficiency. The training curves for both the ProDMM-seq2seq training loss and validation loss are presented in Figure S7. The hyper-parameters configuration are listed in Table A4.

### A.4 Zero-shot Prediction Evaluations of ProDMM-Encoder

#### Perplexity-based Zero-shot Sequences Ranking

Perplexity is a function that maps a sequence to a real-coded value, reflecting the likelihood of the sequence occurring [44]. In the context of Masked Language Models (MLMs) such as BERT [10], the perplexity cannot be directly computed as in traditional autoregressive models [5], since MLMs are bidirectional and designed to predict masked tokens. Thus, we employ the *pseudo-perplexity*, can be calculated for MLMs by iteratively masking and scoring tokens in a given sequence. Specifically, given a sequence **S** = {*w*_1_*, w*_2_*, . . ., w_n_*}, the pseudo-perplexity is computed by masking each token *w_i_* in turn and computing the log-probability assigned by the MLM to the true token *w_i_* given the context **S**_\_*_i_*, where **S**_\_*_i_* denotes the sequence with the *i*-th token masked. Specifically, the contribution of each token to the pseudo-perplexity is given by:

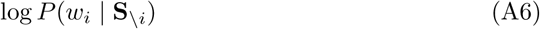

**Table A4:** Pre-training hyper-parameters of ProDMM-Seq2Seq.

| Parameter | Value |
| --- | --- |
| Dropout | 0.1 |
| Hidden size | 1536 |
| Intermediate size | 6144 |
| LayerNorm epsilon | $1.0 \times 10^{-5}$ |
| Number of attention heads | 64 |
| Number of hidden layers | 12 |
| Training mini-batch size | 4096 |
| Evaluation mini-batch size | 4096 |
| Seed | 42 |
| Training steps | 100,000 |
| Gradient Accumulations | 8 |
| Learning rate | $4 \times 10^{-4}$ |
| Adam beta1 | 0.9 |
| Adam beta2 | 0.999 |
| Adam epsilon | 1e-7 |
| Gradient clip value | 1.0 |
| Gradient clip algorithm | norm |
| Precision | BF16 |
| Weight decay | 0.01 |
| Warmup steps | 5000 |

where *P* (*w_i_* | **S**_\_*_i_*) is the probability of token *w_i_* being predicted correctly at the masked position. The overall pseudo-perplexity for the entire sequence **S** is then defined as:

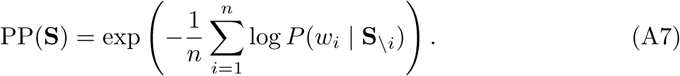

The lower the pseudo-perplexity, the more likely the sequence is under the MLM’s learned distribution. We can utilize the pseudo-perplexity to rank a set of sequences S = {**S**_1_, **S**_2_*, . . .,* **S***_k_*} by computing the pseudo-perplexity for each sequence **S***_j_* and sorting them in ascending order or descending order, depending on the task context.

#### Perplexity-based Zero-shot Mutants Ranking

“Zero-shot” denotes that the model is able to generate prediction results without requiring additional task-specific training or labeled data. Previous studies have demonstrated that when trained on extensive and varied protein sequence databases, protein language models can predict experimental measurements of protein mutant fitness without further supervision [5, 15, 17]. Because ProDMM is trained on both protein and DNA sequences, it can predict the effects of mutants in protein sequences, as well as in CDS and NCDS.

Formally, we denote a *k*-order mutant of a sequence **S** by a set of triplets:

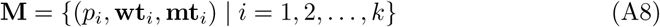

where 1 ≤ *p_i_* ≤ *n* represents the mutant position, *n* is the length of the sequence, **mt***_i_* is the mutant token, and **wt***_i_* is the original token. Specifically, when *k* is 1, the mutant is a single-point mutation. The mutant effect prediction is computed by its log odds ratio:

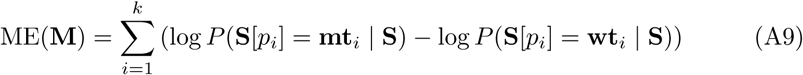

Given a set of mutants of a sequence M = {**M**_1_, **M**_2_*, . . .,* **M***_n_*}, we can use this Equation A9 to rank the mutants based on their predicted effects. Specifically, the mutants with higher ME(**M***_i_*) scores are considered to have a stronger predicted effect, either beneficial or deleterious, depending on the task context.

#### A.4.1 Coding DNA Sequence Expression Prediction

Codon usage bias complexly affects protein expression levels when recombinant proteins are heterologously expressed [45]. Given that ProDMM has been trained on coding DNA sequences, we hypothesize that the expression level of a CDS is negatively correlated with its pseudo-perplexity. This assumption follows that a lower pseudo-perplexity reflects a higher likelihood of the sequence occurring in the learned distribution of the model, indicating that the sequence is more likely to be biologically reasonable and efficient in expression.

##### Dataset

To validate this hypothesis, we first gather a dataset in which [11] experimentally evaluated the expression levels of 6 distinct proteins, each expressed using different coding sequences in the *E.coli* and a real-coded value represents their expression level.

##### Scoring Method

Given a set of CDS sequences that translated into the same protein, we compute the pseudo-perplexity for each CDS according to Equation A7 and rank them in ascending order of their pseudo-perplexity.

##### Evaluation Metrics

We evaluate the ProDMM-Encoder and other models by computing the Spearman ranking correlation coefficient between the model-derived scores and truth expression level. Meanwhile, we utilize a two-tailed t-test to assess whether the performance difference between the ProDMM-Encoder and the previous state-of-the-art model is significant.

#### A.4.2 Zero-shot Non-Coding DNA Sequence Mutant Effect Prediction

Non-coding DNA Sequences are stored in the genome and play crucial roles in protein synthesis and gene regulation. Mutant to NCDS may enhance or decrease its function fitness (Fitness is a metric specific to a study that quantifies how effectively an NCDS performs a particular function). Since ProDMM-Encoder have been trained with Non-coding DNA Sequences, we would like to test its performance on zero-shot NCDS mutant effect prediction.

##### Dataset

We utilize the NCDS mutant effect prediction benchmark from Evo [7]. The benchmark contains 7 NCDS mutant datasets from literature, containing 4 ribozyme mutant datasets [46–49], 2 tRNA mutant datasets [50, 51], and an rRNA mutagenesis dataset [52].

##### Scoring Method

We hypothesize that the mutant effect level of an NCDS is positively correlated with its log odds ratio. We score the mutants according to Equation A9.

##### Evaluation Metrics

We utilize the absolute value of the Spearman ranking correlation coefficient between the model-derived scores and truth mutant fitness. A two-tailed t-test assesses whether the performance difference between the ProDMM-Encoder and the previous state-of-the-art model is significant.

#### A.4.3 Zero-shot Protein Coding DNA Sequence Mutant Effect Prediction

Mutations to a protein coding sequence can affect its various function fitness. Pre-training on prokaryote proteins and coding sequences enables ProDMM-Encoder to perform zero-shot predictions on protein coding sequence mutants [5, 7, 15].

##### Dataset

We gather the protein coding sequence mutant effect prediction benchmark from Evo [7] and ProteinGym [17]. The ProteinGym benchmark contains millions of assay-label protein mutants from various taxons. However, it does not supply the protein coding DNA sequence. Evo [7] collects coding DNA sequence and their codon mutations for six proteins from the ProteinGym benchmark [53–58] that tested in the *E. coli* expression system.

##### Scoring Method

The mutant scoring is also based on the hypothesis that the mutant effect level of a protein-coding sequence is positively correlated with its log odds ratio. The mutant score can be scored with two methods with Equation A9. The first method scores the mutants based on the CDS, while the second is based on the protein sequence. To distinguish the two methods, the first method is denoted as “ProDMM-Encoder (CDS)” and the second is denoted as “ProDMM-Encoder (Protein)”

##### Evaluation Metrics

The scoring results are evaluated with the absolute value of the Spearman ranking correlation coefficient between the model-derived scores and truth mutant fitness.

#### A.4.4 Zero-shot Protein Sequence Mutant Effect Prediction

Protein language models have demonstrated zero-shot prediction ability on protein mutant effect prediction [5, 15].

##### Dataset

As the pre-training dataset of ProDMM-Encoder only consists of prokaryotic organisms, we only evaluate it on the proteins from prokaryotes. We assess the ProDMM-Encoder using a subset of the ProteinGym benchmark [17] that contains 50 deep mutational scanning (DMS) datasets of distinct proteins from prokaryotes. The subset is denoted as “ProteinGym (Pro.)”.

##### Scoring Method

We assume that the mutant effect positively correlates with its log odds ratio. The mutants are scored with Equation A9, where the mutant token is the mutant amino acid, the original token is the wild-type amino acid, and the sequence **S** is its wild-type sequences.

##### Evaluation Metrics

We consider each DMS dataset as an individual prediction dataset, where we score each mutant and compute the Spearman rank correlation between the predicted scores and the experimental fitnesses.

#### A.4.5 Zero-shot Promoter Ranking

Promoters, as non-coding DNA sequences, play a crucial role in gene expression and protein synthetic biology. They are key elements that regulate the initiation of transcription, thereby influencing the levels of gene expression. The interaction between non-coding DNA sequences and downstream coding sequences is a product of co-evolutionary processes that influence both DNA transcription and protein translation efficiencies [59]. This interplay underscores the importance of evaluating the strength of a promoter in conjunction with the downstream protein it regulates. ProDMM is pre-trained on the NCDS-CDS-NCDS and NCDS-protein-NCDS sequence; we can compute the pseudo-perplexity of the promoter as a proxy of expression level.

##### Dataset

We obtained the dataset from [60], which includes various combinations of different promoters and enhanced green fluorescent protein (EGFP, NCBI ID: AAB02572.1) along with their corresponding expression levels.

##### Scoring Method

We assume the conditioned pseudo-perplexity of the promoter is positively correlated with the expression level. The conditioned pseudo-perplexity is:

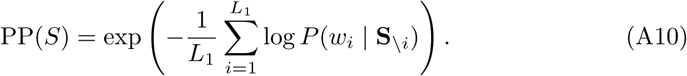

where **S** is the concatenation of the NCDS sequence and CDS (protein) sequence. **S** = {*q*_1_*, q*_2_*, . . ., q_L_*_1_ *, c*_1_*, c*_2_*, . . ., c_L_*_2_ }, in which *q_i_* is promoter (NCDS) token and *c_i_* is CDS (protein) toke as well as *L*_1_ and *L*_2_ are the length of promoter and CDS (protein) respectively.

##### Evaluation Metrics

We utilize Equation A10 to score each combination sequence of promoter and CDS and we use the Spearman rank correlation between the score predicted by the model and the ground truth expression level.

### A.5 Supervised Fine-tuning ProDMM-Encoder on Down-stream Tasks

#### A.5.1 Fine-tuning configurations

During the fine-tuning process, CDS or protein sequences are input into the frozen, pre-trained ProDMM-Encoder, and the mean value of each output serves as the proxy for each sequence. These proxies are then fed into task-specific downstream heads. These heads are all composed of two multiple-layer perceptions (MLPs), with two dropout layers and one GELU activation function, but their parameters vary based on the specific supervised signals used for fine-tuning. All the results are calculated from fivefold cross-validation.

#### A.5.2 Tasks

The ProDMM-Encoder is assessed across four downstream tasks with inputs comprising either protein sequences or CDS sequences. These tasks include predicting the melting point, solubility, protein abundance, and transcript abundance of various native protein sequences. For the transcript abundance predictions, the model’s performance is evaluated on two species: Escherichia coli (E. coli) and Haloferax volcanii (H. volcanii).

#### A.5.3 Dataset

All datasets used in the fine-tuning process are downloaded from the https://github. com/oxpig/CaLM. Five-fold validation is performed using the “KFold” function from the scikit-learn package.

#### A.5.4 Evaluation Metric

Pearson’s correlation coefficient is used as the evaluation metric, which is consistent with CaLM [23]. Pearson’s R is calculated as follows:

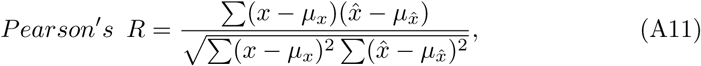

where x and *x̂* represent truth values and predicted values, with *µ_x_* and *µ_x̂_* being their respective means. The calculation is performed using the ”pearsonr” function from the SciPy library.

#### A.5.5 Baselines

The fine-tuned ProDMM-Encoder is benchmarked against 17 models, categorized into two types. The first type are the models that utilize only protein sequences: ESM2-6 [5], ESM2-12, ESM1-6 [61], ESM1-12, ESM1b [24], ESM2-30, ESM2-33, ProTrans BERT (BFD) [6], ProtTrans BERT, ESM2-36, ProTrans T5 XL, ProTrans T5 XL (BFD), ProTrans T5 XXL, ProTrans T5 XXL (BFD), ESM2-48, and ESM1v [15]. The second type is CaLM [23], which received only coding sequences. To ensure a fair comparison, the ProDMM-Encoder is tested on two different input types: protein sequences and CDS, labeled ProDMM pro and ProDMM cds, respectively.

### A.6 ProDMM-Seq2seq

#### A.6.1 Protein Reverse Translation

##### Task

During the fine-tuning phase, we froze the encoder of ProDMM and trained only the decoder. The input sequence to the encoder consisted of the protein sequence appended with the *<*aas2cds*>* prompt token, and the decoder outputed the corresponding CDS.

##### Dataset

We obtained the dataset from [25], which includes various E. coli and human high-expression gene data including protein sequence and CDS. Due to the limited availability of high-expression E. coli data, we also collected non-high-expression gene data to assist in pre-training and fine-tuning our analysis (Figure S4). After applying the Amino Acid–Codon Mask, we observed that even without utilizing the pre-training and fine-tuning pipeline, the performance remained exceptional.

##### Evaluation Metrics

To evaluate the performance of the trained models, we compared both accuracy and perplexity (PPL). Accuracy was measured by generating a reverse-translated DNA sequence from a given protein and then assessing how many codons were predicted correctly. A lower perplexity in our transducer model indicates reduced uncertainty over the distribution of codons. For neural models, perplexity was calculated by exponentiating the average cross-entropy loss, while for the frequency-based baseline models, it was computed by exponentiating the average negative log-likelihood.

##### Baseline

On E. coli and human benchmarks, we compared the fine-tuned ProDMM with frequency-based methods, including unigram, trigram, and 5-gram models, as well as their ensemble (1,3,5-gram) [26], CD-GPT [27], and the state-of-the-art method Transducer [25]. For the zero-shot setting without fine-tuning, ProDMM was compared with CD-GPT, which was also pre-trained with reverse translation capabilities.

#### A.6.2 Co-evolving Effect

##### Task

During the sequence-to-expression fine-tuning phase, we utilized the Encoder of ProDMM augmented with a linear layer as the head. During training, we froze the Encoder and trained only the head. The input sequence is represented in the form ‘N‘.

##### Dataset

We collected coding sequences (CDS) of *E. coli* and their adjacent non-coding regions, along with the corresponding expression levels, from DeepExpression and Ensemble databases. The input sequence comprises the CDS and 300 bp upstream and downstream non-coding regions, which include promoters, 5’ UTRs, 3’ UTRs, and terminators.

##### Evaluation Metric

The evaluation metric is the coefficient of determination (*R*^2^), calculated using the following formula:

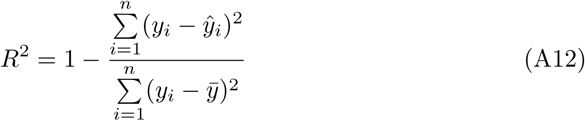

In this formula, *y_i_*represents the true expression value, *ŷ_i_* is the predicted expression value, *ȳ* denotes the mean of the true expression values, and *n* is the number of samples.

#### A.6.3 CDS-guided Promoter Generation

##### Task

During the fine-tuning phase, we froze the encoder of ProDMM and trained only the decoder. The input to the encoder is the CDS sequence appended with the *<*cds2pre-ncds*>* prompt token, and the decoder outputs the corresponding promoter sequence.

##### Dataset

We collected CDS and their corresponding promoter data of E. coli from RegulonDB [31] to construct a benchmark for CDS-to-promoter generation.

##### Baseline

As we are the first to study generating specific promoters from CDS, the available baselines are limited. Therefore, we designed the CDS-Pro Transducer based on the Transducer model [25] as a baseline, and also extended Evo [7] for CDS-NCDS cross-modal generation.

#### A.6.4 Promoter-specific CDS Generation

##### Task

During the fine-tuning phase, we froze the encoder of ProDMM and trained only the decoder. The input to the encoder is the CDS sequence, and the decoder outputs the corresponding promoter sequence.

##### Dataset

We collected CDS and their corresponding promoter data of E. coli from RegulonDB [31] to construct a benchmark for promoter-to-CDS generation.

##### Baseline

As we are the first to study generating specific CDS from the promoter sequence, the available baselines are limited. Therefore, we designed the Pro-CDS Transducer based on the Transducer model [25] as a baseline, and also extended Evo [7] for NCDS-CDS cross-modal generation.

